# Differential amplicons for the evaluation of RNA integrity extracted from complex environmental samples

**DOI:** 10.1101/401109

**Authors:** Fabien Cholet, Umer Z. Ijaz, Cindy J. Smith

**Affiliations:** Infrastructure and Environment Research Division, School of Engineering, University of Glasgow, UK, G12 8LT

**Keywords:** RNA, mRNA, RNA integrity, RIN, differential amplicons, R_amp_

## Abstract

**Background:** Reliability and reproducibility of transcriptomics-based studies are highly dependent on the integrity of RNA. Microfluidics-based techniques based on ribosomal RNA such as the RNA Integrity Number (RIN) are currently the only approaches to evaluate RNA integrity. However, it is not known if ribosomal RNA reflects the integrity of the meaningful part of the sample, the mRNA. Here we test this assumption and present a new integrity index, the Ratio amplicon, R_amp_, to monitor mRNA integrity based on the differential amplification of long to short RT-Q-PCR amplicons of the glutamine synthetase A (*glnA*) transcript.

**Results:** We successfully designed and tested two R_amp_ indexes targeting *glnA* transcripts. We showed in a suite of experimental degradations of RNA extracted from sediment that while the RIN in general did reflect the degradation status of the RNA well the R_amp_ mapped mRNA degradation better as reflected by changes in Reverse Transcriptase Quantitative PCR (RT-Q-PCR) results. Furthermore, we examined the effect of degradation on transcript community structure by amplicon sequencing of the *16S rRNA, amoA* and *glnA* transcript which was successful even form the highly-degraded samples. While RNA degradation changed the community structure of the mRNA profiles, no changes were observed between successively degraded 16S rRNA transcripts profiles.

**Conclusion:** As demonstrated, transcripts can be quantified and sequenced even from highly degraded samples. Therefore, we strongly recommend that a quality check of RNA is conducted to ensure validity of results. For this both the RIN and R_amp_ are useful, with the R_amp_ better evaluating mRNA integrity in this study.

## Background

A key question in environmental microbiology is to determine the functioning and activity of microbial communities. While genomic approaches have resulted in an unprecedented understanding of their structure and complexity [1], they do not inform of the actual activity and functioning at a given time. In this case targeting the transcriptome, that is the subset of genes that are actively transcribed at a given time, is more informative. While there can be substantial post translational regulation that may prevent final protein synthesis and/or activity, gene expression is the direct link between the genome and the function it encodes and therefore is a stronger link to activity than DNA approaches alone [2]. As a result transcriptomics based approaches are widely used to assess microbial activity and functioning in the environment [3, 4]. The premise is that that messenger RNA (mRNA) turn-over within cells is rapid, ranging from a few minutes to less than an hour [5]. As such a snap-shot of the transcriptome reflects the cells transcriptional response to its surrounding environment and metabolic needs at a given time.

A challenge for all transcript-based studies, not least for those from environmental samples, is to ensure the quality and integrity of the RNA on which the results are based. Extracted RNA is prone to degradation both during the extraction procedure, post-extraction handling and over time. Factors such as RNase activity, physical degradation during extraction procedures, and even storage can degrade RNA. If there is significant post-extraction degradation among different samples that are to be compared, the interpretation of results may be compromised. In other words, differences between samples may arise as a result of post-extraction degradation, as opposed to representing actual difference in gene expression. Indeed, meaningful and reproducible results can only be obtained when working with good quality, intact RNA, whether it is eukaryotic RNA [6–9] or Prokaryotic RNA [10]. As such an initial quality check of extracted RNA, not least from complex environmental microbial communities should be the essential first step before proceeding to any downstream applications. This quality check would help to ensure that any differences observed between samples are due to actual changes in gene expression rather than differences in samples integrity as a result of degradation.

Current methods to evaluate the integrity of extracted RNA are based on ribosomal RNA (rRNA). These approaches evaluate integrity as a ratio between the 16S and 23S ribosomal RNA: 16S, 23S and 5S rRNA are synthetized as one primary transcript and are separated upon maturation [11]. The 16S and 23S ribosome should therefore be present at a ratio 1:1. However, as the 23S ribosome is approximately twice as large as the 16S ribosome, for intact, non-degraded RNA, the expected ratio of 23S:16S RNA is 2:1. However, the caveat of this approach is the assumption that the integrity of rRNA reflects that of the overall RNA, including mRNA. The relationship between the integrity of rRNA and that of mRNA has not been demonstrated [6]. Indeed, there are several reports indicating the more stable properties of rRNA compared to mRNA [12–14]. As such, the usefulness of this ratio to assess mRNA integrity is still unclear.

In its simplest form, evaluating ribosomal RNA integrity is an electrophoretic separation of RNA in a gel matrix. Essentially, a visual check for the presence of the characteristic bands corresponding to 16S and 23S rRNA. More advanced techniques based on microfluidics are better suited for assessing RNA quality, allowing for the calculation of integrity indexes, such as the RNA Integrity Number, RIN (Agilent Technologies) or the RNA Quality Score, RQI (BioRad). These scores vary between 0 (RNA totally degraded) and 10 (“perfect” RNA). A value of 7 has been suggested as a limit between “good” and “bad” quality RNA extracted from bacterial pure cultures [10]. However, RNA extracted from natural environments such as soil or sediment will likely have lower quality due to the more complex matrixes and, often, harsh extraction techniques routinely used, such as bead beating [15] but this information is not widely reported in the literature. Nevertheless, as highlighted above, even if reported, a shortcoming for RIN/RQI algorithms is that they are primarily based on rRNA (16S/23S ratio) which may degrade differently from mRNAs; the relevance of such indexes for gene expression analysis is therefore unknown.

In Eukaryotic gene expression studies, an alternative index often used to evaluate the level of mRNA degradation is the 3’-5’ ratio [16]. This technique is based on the observation that Eukaryotic mRNAs generally degrade from the 5’ to the 3’ end, with the 3’ poly A tail acting as a protective agent. As a result, Reverse Transcriptase-PCR (RT-PCR) targeting the 5’ end of the transcript is less likely to be produce amplicons than those targeting the 3’ end. A high 3’:5’ ratio (low 5’ copy number) is therefore an indication of mRNA degradation. This technique cannot be applied to prokaryotic mRNAs as they generally don’t possess poly A tails, and when they do, the tail enhances mRNA degradation [17]. Recently, a new approach called differential amplicon (Δamp) has been developed [18]. This technique is based on the differential amplification of RT-PCR amplicons of different lengths from the same mRNA target as a new means to determining RNA integrity (see also [19]). Here it was observed that the copy number of long RT-Q-PCR targets correlated with mRNA degradation whereas the short targets were more stable. Since this approach doesn’t rely on the presence of the poly A tail, it could theoretically be adapted to prokaryotic mRNAs. Although this has not been directly observed for prokaryote RNA, Reck *et al* [20] showed a similar response of a exogenous green-fluorescent-protein mRNA(GFP) that they spiked into stool RNA to montior its intergery when subjected to different storage conditions. They showed that the copy number of the spiked exogenous GFP correlated well with RNA integrity when targeting a long amplicon (≥500bp), whereas the short amplicon (≤100bp) remained constant, even in highly degraded RNA preparations. This indicated that, as was observed by Björkman and co-workers, longer mRNA targets reflect degradation better. As such, the difference in RT-Q-PCR performance, reflected by the difference in cycle threshold (Ct) between a short and a long amplicon from the same cDNA target could be used as an index to reflect mRNA integrity.

Here, we propose to exploit the differential amplicon approach to develop a ratio of long to short amplicons directly targeting mRNA transcripts of the same target but of differing lengths by RT-Q-PCR as an indicator of overall mRNA integrity. For this we propose the ubiquitous bacterial glutamine synthetase A transcript (*glnA*) as the target. Glutamine synthetase is a ubiquitous gene, found in Bacteria and Archaea [21, 22], with a role in assimilating inorganic nitrogen (ammonia) into amino acids [23]. The *glnA* transcript has been used previously in RT-(Q)-PCR approaches to evaluate RNA extraction yield from soils [24–26]. However, as the expression of *glnA* is regulated by ammonia concentration [27–29], the copy number of this transcript can vary making comparison between studies difficult. Our approach overcomes this difficulty by calculating the ratio of long to short *glnA* transcripts. We designate this the Ratio Amplicon (R_amp_), and propose it as an indicator of mRNA integrity, independent of absolute gene expression.

Specifically, this study aims to design and test the Ratio Amplicon (R_amp_) approach to evaluate bacterial mRNA integrity extracted from marine sediments. Furthermore, we aim to compare and evaluate this approach against the conventional ribosomal based RNA integrity index, RIN. Comparison between the two approaches was conducted by monitoring how well both indexes reflected experimental RNA degradation (UV, heat, RNase, freeze/thaw). The impact of RNA degradation and the ability of the two indexes to predict ribosomal and mRNA integrity was evaluated via quantification of two commonly surveyed bacterial transcripts, the highly abundant ribosomal 16S rRNA and mRNA from the less abundant bacterial ammonia monooxygenase (*amoA*). Finally, the effect of RNA degradation on transcript community structure was evaluated by amplicon sequencing of the cDNA obtained from sequentially degraded samples.

We hypothesised that i) the R_amp_ would be a better predictor of mRNA integrity than the RIN and ii) RNA degradation would adversely affect both transcript quantification and community composition.

## Methods

### Sediment Samples

Surface mud samples (0 to 2 cm) were collected on 11/01/2017 from Rusheen Bay, Ireland (53.2589° N, 9.1203° W) (presence of *amoA* genes/transcripts previously established [30, 31] in sterile 50ml Eppendorf tubes, flash frozen and stored at -80°C until subsequent use.

### Design of new *glnA* primers

To design new primers, bacterial *glnA* sequences were downloaded from the GeneBank database [32]. Sequences related to environmental bacteria were subjected to BLAST search [33] in order to gather additional sequences. In total eighty-four sequences (Additional file 1) were aligned using MUSCLE [34] and a phylogenetic neighbour joining tree was drawn in MEGA 7 [35]. Based on sequence similarity, eight groups could be distinguished (see Additional file 4: Figure S1). Primer sequences from Hurt and co-workers [15] were aligned in each individual group to determine coverage and new primers (Table 1) were designed based on conserved regions to target the same groups with varying length primers.

**Table 1.**
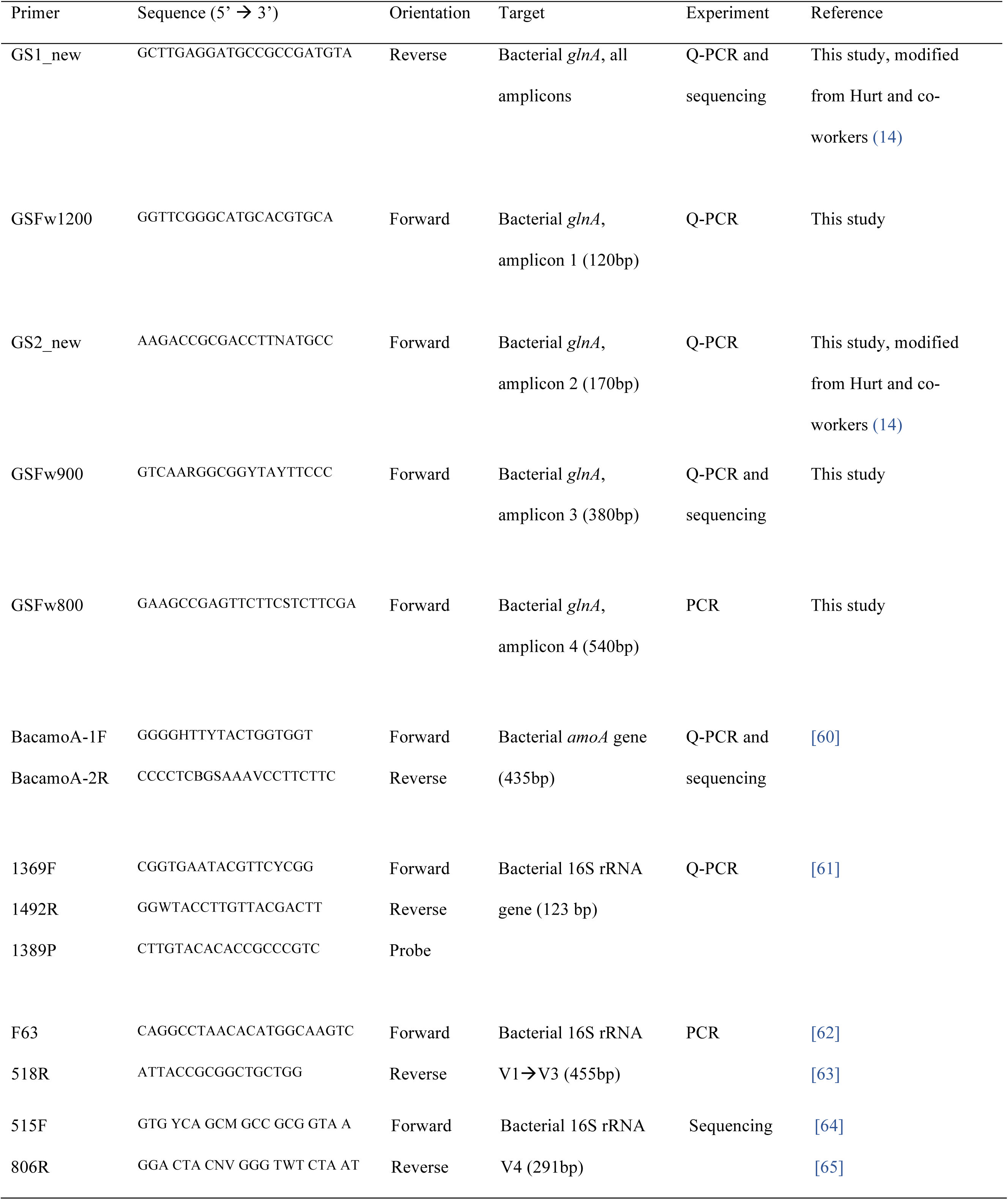
List of primers used in this study.

Primers were tested on DNA and cDNA using environmental DNA/RNA extractions and environmental cDNA, as template. *glnA* genes were amplified (BIOTAQ DNA polymerase kit; Bioline) in a 25µl final volume composed of 2.5µl BioTaq10x buffer, 18µl water, 1.5µl MgCl_2_ (50mM), 0.5µl of each primer (10µM), 0.5µl dNTPs (10µM each), 0.5µl *Taq* DNA polymerase and 1µl of template. PCR conditions were as follow: 95°C 5 min, (94°C 30 sec, 60°C 30 sec, 72°C 30 sec) x 30 and 72°C 5 min.

### RNA preparation from sediment

All surfaces and equipment were cleaned with 70% ethanol and RNase Zap (Ambion) before sample processing. All glassware and stirrers used for solutions preparation were baked at 180°C overnight to inactivate RNases. All plasticware was soaked overnight in RNase away (ThermoFisher Scientific) solution. Consumables used, including tubes and pipet tips were RNase free. All solutions were prepared using Diethylpyrocarbonate (DEPC) treated Milli-Q water. A simultaneous DNA/RNA extraction method, based on that of Griffiths and co-workers [36] was used to recover nucleic acids from sediment. Briefly, 0.5g of sediments were extracted from using bead beating lysing tubes (Matrix tube E; MP Biomedical) and homogeneised in 0.5ml CTAB/phosphate buffer (composition for 120 ml: 2.58g K_2_HPO_4_.3H_2_O; 0.10g KH_2_PO_4_; 5.0g CTAB; 2.05g NaCl) plus 0.5ml Phenol:Chlorophorm:Isoamyl alcohol (25:24:1 v:v:v). Lysis was carried out on a FastPrep system (MP Biomedical) (S: 6.0; 40sec) followed by a centrifugation at 12,000g for 20min (4°C). The top aqueous layer was transferred in a fresh 1.5ml tube and mixed with 0.5ml chlorophorm:isoamyl alcohol (24:1 v:v). The mixture was centrifuged at 16,000g for 5min (4°C) and the top aqueous layer was transferred in a new 1.5ml tube. Nucleic acids were precipitated by adding two volumes of a solution containing 30% poly(etlyleneglycol)_6000_ (PEG6000) and 1.6M NaCl for 2 hours on ice and recovered by centrifugation at 16,000 x g for 30 min (4°C). The pellet was washed with 1ml ice-cold 70% ethanol and centrifuged at 16,000g for 30 min (4°C). Ethanol wash was discarded and the pellet was air dried. Once the ethanol was completely evaporated, the pellet was re-suspended in DEPC treated water. DNA/RNA preparations were stored at -80°C if not used immediately.

RNA was prepared from the DNA/RNA co-extraction by DNase treating with Turbo DNase Kit (Ambion) using the extended protocol: half the recommended DNase volume is added to the sample and incubated for 30min at 37°C, afterwhich the second half of DNase is then added and the sample is re-incubated at 37°C for 1 hour. Success of the DNase treatment was checked by no PCR amplification of the V1-V3 Bacterial *16S rRNA* gene [4].

### RNA degradation experiments

#### Physical degradation

To obtain RNA with controlled degradation status, DNA free RNA preparations (≈8µl) were aliquoted from an initial extraction in separate 0.2ml RNase free tubes and incubated at 90°C or under a UV lamp for 0, 10, 45 or 90 minutes. To determine the potential effect of repeated freeze-thaw on RNA preparations, the same 15µl DNA-free RNA was exposed to cycles of freezing (at -80°) and thawing (on ice) as follows - 0, 1, 3, 5, 7 and 10 freeze-thaw cycles. cDNA was then generated for each individual aliquot as described later.

#### Enzymatic Degradation by RNase I

For RNAse I degradation experiment, 40µl aliquots of DNA-free RNA was incubated at 37°C for 40min in the presence of increasing concentrations of RNaseI (supplier): 0 (buffer only), 2, 10, 20 and 40 Units RNase I/ µg RNA. The reaction was stopped by adding 10µl β-mercaptoethanol and RNA was recovered by ethanol precipitation: 5µl of 7.5M ammonium acetate and 137.5µl 100% ethanol was added and the mixture was precipitated overnight at - 20°C. RNA was pelleted by centrifugation 16,000 x g for 40min at 4°C and the pellet was washed with 480µl ice cold 70% ethanol and pelleted by centrifugation at 16,000g for 30min at 4°C. The pellet was air dried and re-suspended in 40µl of DEPC-treated water. An aliquot of RNA that did not undergo ethanol precipitation was also included for comparison (designated NT: “Not Treated”).

### Reverse Transcriptase Reaction

DNA-free RNA was used for *glnA* cDNA synthesis using Superscript III kit (Invitrogen) and gene specific priming. The initial RT mixture containing 3µl water, 1µl reverse primer GS1_new (10µM), 1µl dNTP’s (10mM each) and 5µl template was incubated at 65°C for 5 min and quickly transferred on ice for 1 min. A second mix composed of 4 µl 5X first-strand buffer, 1 µl 0.1 mM dithiothreitol (DTT), and 1µl SuperScript III (200 units/µl) was added and the resulting mixture was incubated at 55°C for 50 min and then at 72°C for 15 min. The primers and PCR conditions for the amplification of *glnA* targets from cDNA were similar to those used for DNA.

For *16S rRNA* and *amoA* genes, Superscript III kit (Invitrogen) and random hexamers priming was used. The initial RT mixture containing 3µl water, 1µl random hexamer (50µM), 1µl dNTP’s (10mM each) and 5µl template was incubated at 65°C for 5 min and quickly transferred to ice for 1 min. A second mix composed of 4 µl 5X first-strand buffer, 1 µl 0.1 mM dithiothreitol (DTT), and 1µl SuperScript III (200 units/µl) and 1µl RNase inhibitor (40U/µl) was added and the resulting mixture was incubated at 25°C for 5 min, 55°C for 50 min and then at 72°C for 15 min.

### RNA integrity evaluation

#### RNA integrity number

RINs were determined at all degradation points, using the automated 2100 Bioanalyser platform (Agilent Technologies) with the Prokaryote total RNA Nano chip, following the manufacturer’s instructions.

#### glnA Q-PCR and Ratio amp (R_amp_) calculation

*glnA* cDNA underwent Q-PCR, to amplify varying length amplicon fragments with primer combination as detailed in table 1). Three *glnA* amplicons were produced (Fig. 1), a 120bp amplicon (amplicon 1) generated using the primer pair GS1_new/GSFw1200, a 170bp amplicon (amplicon 2) generated using the primer pair GS1_new/GS2_new and a 380bp amplicon (amplicon 3) generated using the primer pair GS1_new/GSFw900. Q-PCR reaction (10µl) was composed of 5µl EVAGreen Supermixes (SsoFast; Bio-Rad), 0.3µl of each primers (10µM) and 1µl of cDNA template (1/10 diluted). The Q-PCR condition was as follows: 95°C-30sec, (95°C-10sec; 65°C-10 sec) x 35 cycles; plate read at 65°C. Melt curve analysis was performed from 65°C to 95°C with 0.5°C increment every 5 sec.

**Fig 1.**
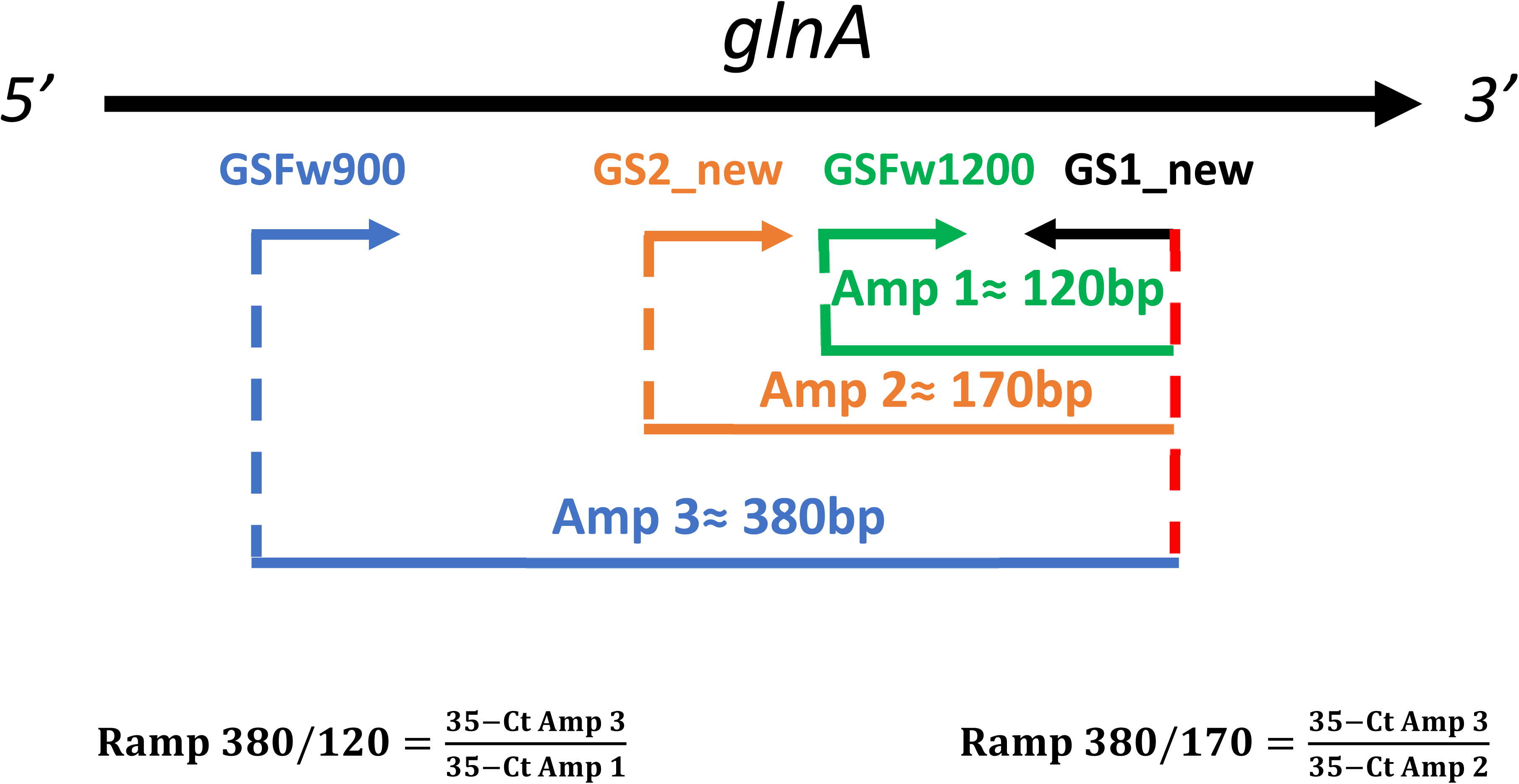
Schematic representation of primer binding sites along the Bacterial *glnA* gene. Primers are represented by arrows pointing to the right (forward primers) or to the left (reverse primer). The amplicons (Amp) generated by the different primer combinations are represented as colored lines. The formulas used to calculate the two R_amp_ indexes are detailed under the figure.

The Ct value of each assay was recorded and the differential amplicon ratios (R_amp_) were calculated for each degradation point as follows:

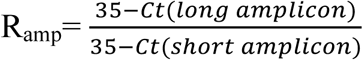

The value of 35 was chosen as the maximum number of Q-PCR cycles the reaction underwent. A transformation of the differential amplicon was applied in order to have a theoretical maximal value of 1 (no degradation of RNA) and a theoretical minimal value close to 0 (totally degraded RNA).

## *amoA* and *16S rRNA* RT-Q-PCR

For all degradation experiments, the Cts of the Bacterial *amoA* and the Bacterial *16S rRNA* was determined by Q-PCR of the cDNA preparations. The *amoA* Q-PCR was carried out in a 20µl reaction volume composed of 10µl 5µl EVAGreen Supermixes (SsoFast; Bio-Rad), 0.4µl of each primer (BacamoA-1F and BacamoA-2R) (10µM each), 7.2µl water and 2µl of cDNA template (1/10 diluted). The Q-PCR cycle was as follows: 95°C-5 min, (95°C-30sec, 47°C-30 sec, 72°C-1min, 81°C-1sec→ plate read) x 40 cycles. Melt curve analysis was performed from 65°C to 95°C with 0.5°C increment every 5 sec. *16S rRNA* cDNA targets were quantified in a 20µl reaction volume composed of 10µl Itaq Universal Probes Supermix (Bio-Rad), 1.8µl each primer (1369F and 1492r) (10µM each), 0.4µl probe (1389P) (10µM), 5µl water and 1µl cDNA template (1/10 diluted). The Q-PCR cycle was as follows: 95°C-10min, (95°C-10sec, 60°C-30sec) x 40 cycles and 40°C-10min. All primers are detailed in table 1.

## Illumina sequencing

The qualitative effect of RNA degradation the community composition of the three bacterial genes (*amoA*, *glnA* and 16S rRNA) was determined by sequencing the amplicons generated from the cDNA preparations obtained after RNAse I degradation. For each PCR amplification was carried out using the HotStartTaq PCR kit (Qiagen) in the following mix 25µl volume: 19.8µl water, 0.5µl of each primer (10µM each), 0.5µl dNTPs (10µM each), 0.2µl HotStartTaq, 2.5µl of 10x PCR buffer and 1µl cDNA template (10^-1^ and 10^-3^ diluted for functional genes and 16S rRNA respectively). Primers used for sequencing are listed in table 1 (Illumina adaptors were added at the 5’ end of the sequencing primers for PCR: 5’-TCG TCG GCA GCG TCA GAT GTG TAT AAG AGA CAG (forward adaptor); 5’-GTC TCG TGG GCT CGG AGA TGT GTA TAA GAG ACA G (reverse adaptor). The PCR cycles were as follows: *amoA*: 95°C-15min, (94°C-30sec, 55°C-30sec, 72°C-30sec) x 32 cycles and 72°C-10min final extension; *glnA*: 95°C-15min, (94°C-30sec, 55.6°C-40sec, 72°C-40sec) x 32 cycles and 72°C-7min final extension; 16S rRNA: 95°C-15min, (94°C-45sec, 50°C-30sec, 72°C-40sec) x 25 cycles and 72°C-10min final extension. For each functional gene, three separate PCRs were carried out, using the same conditions, and pooled together for further processing.

PCR amplicons were cleaned using the AMPure XP beads kit following the manufacturer’s recommendations. Illumina indexes were then attached using the Nextera XT Index Kit with the following PCR condition: 95°C-15min, (95°C-30sec, 55°C-30sec, 72°C-30sec) x 8 cycles and 72°C-5min. The resulting amplicons were purified using the AMPure XP beads kit and eluted in 25µl water. After this step, some preparations were randomly chosen (2 per genes) and run on the Bioanalyser following the DNA 1000 Assay protocol (Agilent Technologies) to determine the average length of the amplicons and to check for the presence of unspecific products. Finally, DNA concentration was determined using fluorometric quantification method (Qubit) and molarity was calculated using the following equation:

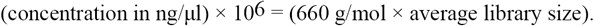

Libraries were pooled in equimolar amount, and checked again on the Bioanalyser and the final library was sent to the Earlham Institute (Norwich Research Park, Norwich, UK) for Illumina MiSeq amplicon sequencing.

## Bioinformatics

### Construction of the reference databases

The following sequences were downloaded (see Additional file 2): *amoA* sequences from Fungene (http://fungene.cme.msu.edu/) alongside NCBI sequences (n=642); and bacterial *glnA* sequences (n=1330) as FASTA files from Microbial Genome Database (http://mbgd.genome.ad.jp). For *amoA s*equences, the NCBI taxonomy was given in the FASTA headers whereas for *glnA* sequences, the MBGD Archive (http://mbgd.genome.ad.jp/htbin/view_arch.cgi) was used to download annotations (mbgd_2016_01) associated with the sequences, and a custom script was written to identify and tag the sequences with NCBI taxonomy. Subsequently, R’s rentrez [37] package was used to get taxonomic information at different levels to generate a taxonomy file for *glnA* sequences. The FASTA file and the corresponding taxonomy file was then formatted to work with Qiime. For *16S rRNA* we used the SILVA SSU Ref NR database release v123.

### Processing of amplicon sequences

Abundance tables were obtained by constructing operational taxonomic units (OTUs) as follows. Paired-end reads were trimmed and filtered using Sickle v1.200 [38] by applying a sliding window approach and trimming regions where the average base quality drops below 20. Following this we apply a 10 bp length threshold to discard reads that fall below this length. We then used BayesHammer [39] from the Spades v2.5.0 assembler to error correct the paired-end reads followed by pandaseq v(2.4) with a minimum overlap of 20 bp to assemble the forward and reverse reads into a single sequence. The above choice of software was as a result of author’s recent work [40, 41] where it was shown that the above strategy of read trimming followed by error correction and overlapping reads reduces the substitution rates significantly. After having obtained the consensus sequences from each sample, the VSEARCH (v2.3.4) pipeline (all these steps are documented in https://github.com/torognes/vsearch/wiki/VSEARCH-pipeline) was used for OTU construction. The approach is as follows: the reads are pooled from different samples together and barcodes added to keep an account of the samples these reads originate from. Reads are then de-replicated and sorted by decreasing abundance and singletons discarded. In the next step, the reads are clustered based on 97% similarity, followed by removing clusters that have chimeric models built from more abundant reads (--uchime_denovo option in vsearch). A few chimeras may be missed, especially if they have parents that are absent from the reads or are present with very low abundance. Therefore, in the next step, we use a reference-based chimera filtering step (--uchime_ref option in vsearch) using a gold database (www.mothur.org/w/images/f/f1/Silva.gold.bacteria.zip) for *16S rRNA* sequences, and the above created reference databases for *glnA* and *amoA* genes. The original barcoded reads were matched against clean OTUs with 97% similarity to generate OTU tables (4108, 1691, and 55 OTU sequences for 16SrRNA, *glnA* and *amoA* respectively). The representative OTUs were then taxonomically classified using assign_taxonomy.py script from Qiime [42] against the reference databases. To find the phylogenetic distances between OTUs, we first multi sequence aligned the OTUs against each other using Mafft [43] and then used FastTree v2.1.7 [44] to generate the phylogenetic tree in NEWICK format. Finally, make_otu_table.py from Qiime workflow was employed to combine abundance table with taxonomy information to generate biome file for OTUs.

## Statistical analysis

All statistical analyses were carried out in R. For degradation experiments RIN and R_amp_ values were compared between time points with one-way ANOVA, when the ANOVA test was significant, differences between time points were investigated using Tuckey HSD post-hoc test. For community analysis (including alpha and beta diversity analyses) we have used the vegan package [45]. To find OTUs that are significantly different between multiple conditions (Degradation), DESeqDataSetFromMatrix() function from DESeq2 [46] package with the adjusted p-value significance cut-off of 0.05 and log2 fold change cut-off of 2 was used. Vegan’s adonis() was used for analysis of variance (henceforth referred to as PERMANOVA) using distance matrices (BrayCurtis/Unweighted Unifrac/Weighted Unifrac for gene sequences) i.e., partitioning distance matrices among sources of variation (Degradation). The scripts for above analysis can be found at http://userweb.eng.gla.ac.uk/umer.ijaz/#bioinformatics

## Results

### Design and optimization of *glnA* primers

Three new forward *glnA* primers (GSFw1200, GSFw900 and GSFw800) were designed to target a conserved region in groups 3, 4, 5, 7 and 8 of the *glnA* alignment (Additional file 4: Figure S1) at a position ≈120 bp, ≈380bp and ≈500bp, respectively, in front (closer to the 5’ end of the gene) of an updated reverse primer from Hurt and co-workers named, GS1_new primer. This resulted in three amplicon sizes to derive a ratio amplicon (R_amp_) from (Fig. 1). The newly designed primers (Table 1) were optimised for PCR and RT-PCR resulting in amplicons of the expected size for all primer pairs. Assays were subsequently optimised for SYBR Green Q-PCR. All primers except for GSFw800, producing the 500 bp amplicon were successfully optimised with diagnostic single peak melt curves. As such we proceeded with two R_amp_ ratio primer sets the R_amp_ 380/120 and the R_amp_ 380/170.

### Heat degradation

Incubation of RNA at 90°C had a strong and rapid impact on its integrity with a drop in the RIN from 7.5 to 4.7 after 10min. At this point, the band corresponding to 23S rRNA had almost completely disappeared. Further exposure resulted in more pronounced degradation with accumulation of short RNA fragments and a RIN around 2 for both 45min and 90min exposure (Fig. 2A & 1A). One-way ANOVA revealed significant difference between all time-points, except 45 and 90min. A low and non-significant decrease in both R_amp_ indexes was observed (−0.07 for 380/120 and -0.11 for 380/170) between 0min and 10min (Fig. 2C). This would tend to indicate that the R_amp_ was less sensitive than the RIN for monitoring RNA degradation by heat. However, interestingly the increase in Ct was also not significant for both *amoA* and *16S rRNA* between 0 and 10min (Fig. 2B), showing that the R_amp_ reflected the outcome of the RT-Q-PCR assays better than the RIN. Further exposure to heat induced a more pronounced decrease in both R_amp_ (≈ -0.4 for 380/120 and ≈ -0.3 for 380/170) at 45min compared to 0min. Both R_amp_ indexes reached values around 0.15 at 90min, which mapped well the behaviour of *amoA*, with a sharp increase in the Ct for this transcript between 10 and 45min (≈4cts) and between 45 and 90min (another ≈4cts). The 16S rRNA transcript was also affected but to a smaller extent (increase in Ct of only ≈3ct between 0 and 90min). Yet, in this case too, the increase was quite low between 0 and 10min and sharper between 10-45min and 45-90min.

**Fig 2.**
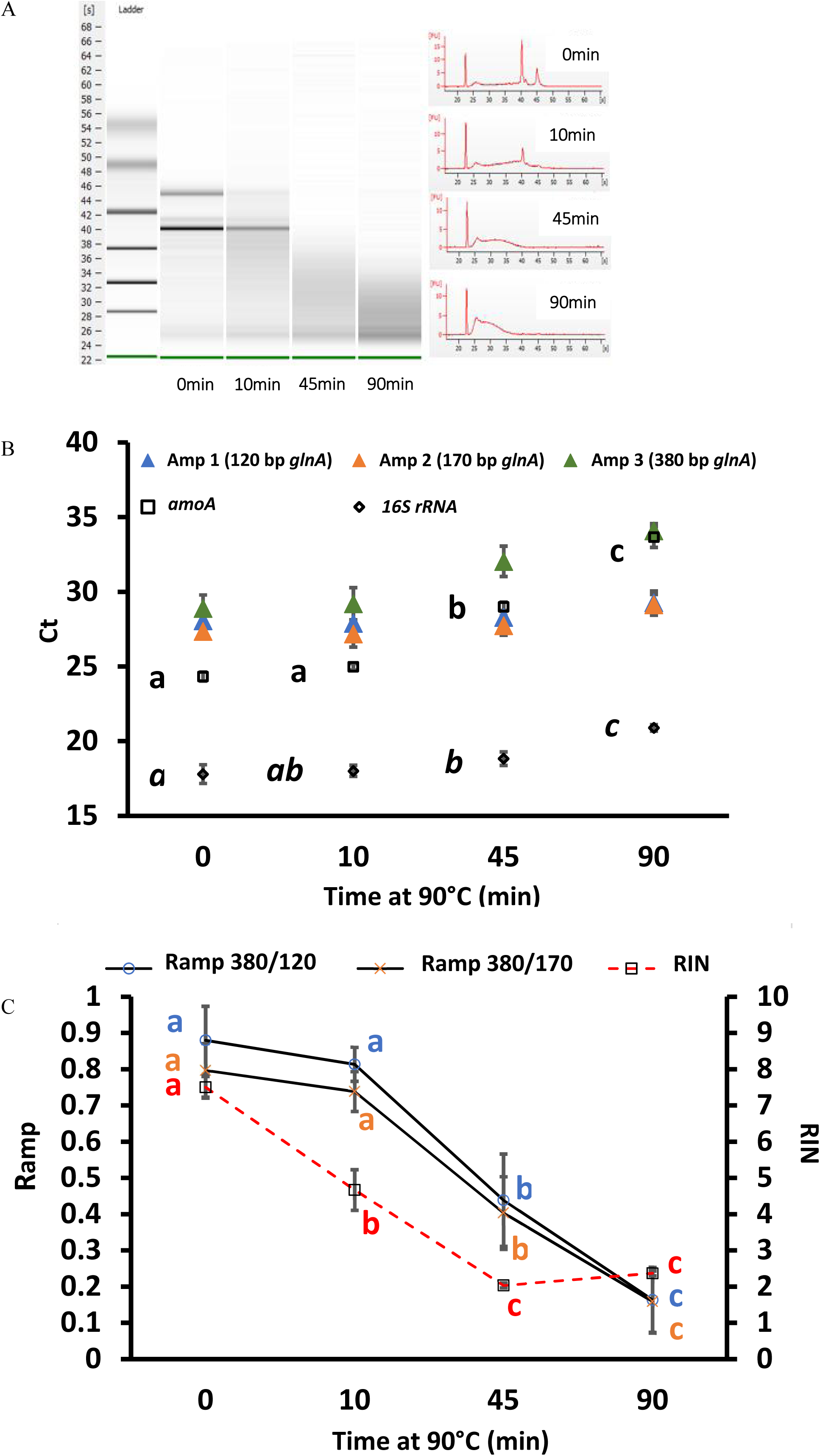
Effect of heat degradation on RNA integrity measured via the RIN (A), with RT-Q-PCR (B) and RIN versus R_amp_ (C). For RIN, RNA integrity visualised in virtual gels (A; left) and electropherogram (A; right) are displayed against incubation period at 90°C. B) Effect of degradation on transcript quantification; Amp 1-3: average Ct (n=3) of one of the three possible *glnA* amplicons; *amoA*: average *amoA* Ct (n=3) of the Bacterial *amoA* transcript; *16S rRNA*: average *16S rRNA* Ct (n=3) of the bacterial *16S rRNA* transcript. Letters indicate the result of TukeyHSD tests (points with different letters had values significantly different from each other using 0.05 as threshold for the *p*.value). Effect of RNA degradation on R_amp_ index is presented in Fig. C. The R_amp_ 380/120 was calculated as 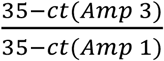 and the Ramp 380/170 as 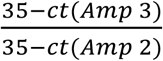; For comparison, RIN values were also plotted.

### UV degradation

The RIN was almost insensitive to UV radiation with an overall decrease of ≈1 at 90min compared to 0min (Fig. 3A & 3C). In contrast, UV radiation had a more pronounced effect on transcript quantification than heat as reflected by a quasi linear increase in Ct of the *amoA* transcript between 0 and 45min (Fig. 3B). Unlike heat exposure, 10min under UV induced strong and significant increase in *amoA* Ct values (≈4cts). At 45min, the Ct had increased by ≈9 compared to the starting point. After 90min, the Ct of the *amoA* transcript almost reached 35, close to the detection limit. The Ct for *16S rRNA* transcript increased steadily from 18 at 0min to 20 at 90min, showing that this assay/transcript was less sensitive to UV degradation. The behaviour of the R_amp_, again, mapped well onto *amoA* behaviour with a decrease of ≈0.2 after 10min exposure for both indexes (though this was not significant) (Fig. 3C). A net decrease was observed at 45min (≈ -0.6 compared to 0min) and at 90min both R_amp_ almost reached 0 since the Ct of the amplicon 3 *glnA* (380bp) was very close to 35.

**Fig 3.**
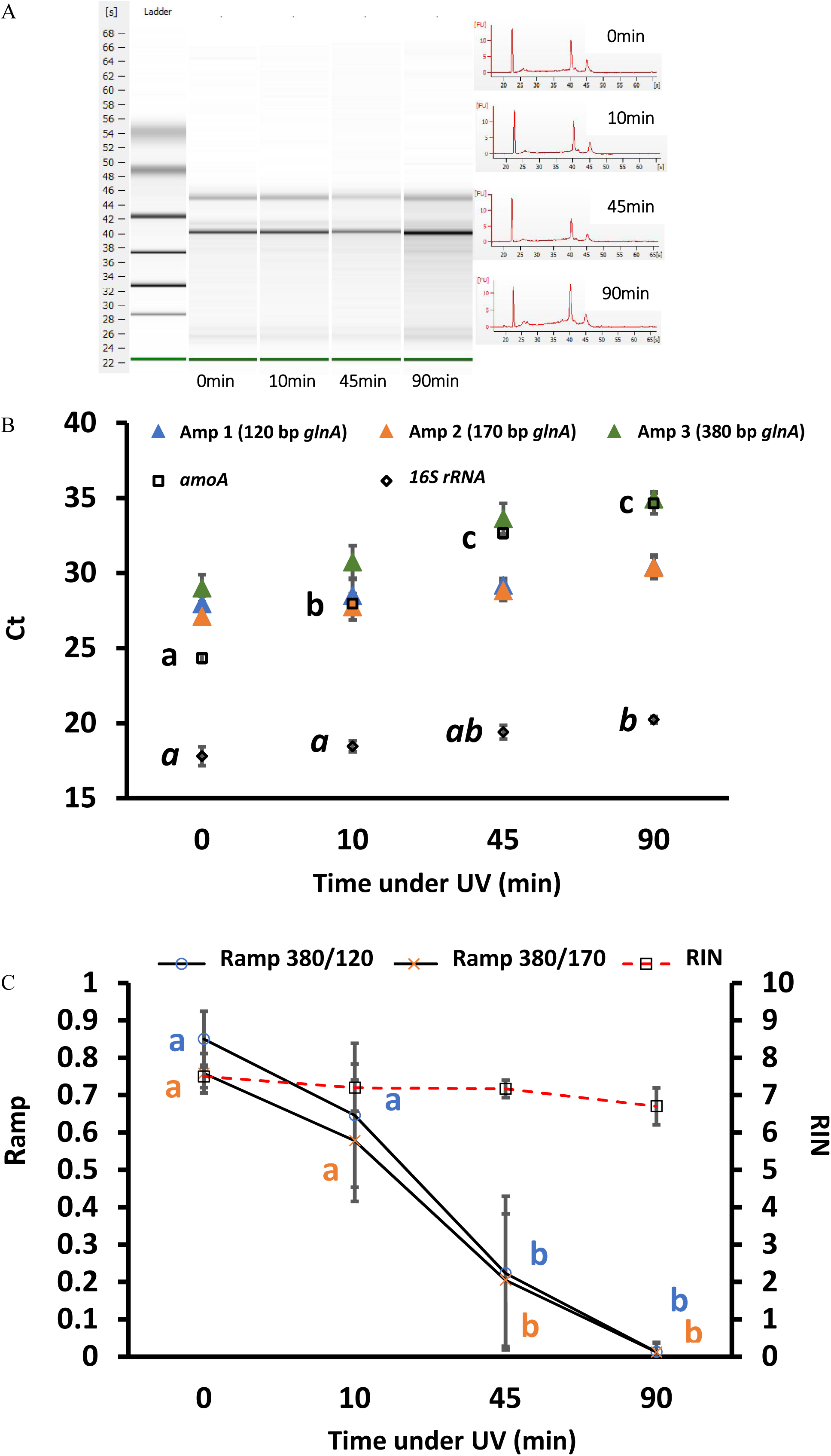
Effect of UV degradation on RNA integrity measured via the RIN (A), with RT-Q-PCR (B) and RIN versus R_amp_ (C). For RIN, RNA integrity visualised in virtual gels (A; left) and electropherogram (A; right) are displayed against incubation period under UV. B) Effect of degradation on transcript quantification; Amp 1-3: average Ct (n=3) of one of the three possible *glnA* amplicons; *amoA*: average *amoA* Ct (n=3) of the Bacterial *amoA* transcript; *16S rRNA*: average *16S rRNA* Ct (n=3) of the bacterial *16S rRNA* transcript. Letters indicate the result of TukeyHSD tests (points with different letters had values significantly different from each other using 0.05 as threshold for the *p*.value). Effect of RNA degradation on R_amp_ index is presented in Fig. C. The R_amp_ 380/120 was calculated as 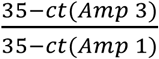 and the Ramp 380/170 as 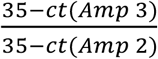; For comparison, RIN values were also plotted.

### Degradation by RNaseI

The RIN showed a rapid response to RNase I degradation with a decrease from 7.1 to 6 between 0 and 2U/µg (Fig. 4A & 4B.) as seen on virtual gels and electropherograms with an almost complete disappearance of the 23S rRNA. When using 10U/µg and higher concentrations, the RIN decreased and remained stable at approximately 2.5 indicating advanced/almost complete degradation of the RNA. Complete destruction of both rRNA and an accumulation of small size RNA molecules on the electropherogram can be observed (Fig 4A). In contrast, enzymatic degradation by RNase I had a relatively small effect on the Ct of the *amoA* transcript at low concentration (only 0.2 Ct increase between 0U/µg and 2U/µg treatments) (Fig. 4B). Ct values for *amoA* increased with greater degradation of the parent RNA (3 Cts difference at 10 and 20U/µg and 5 Cts at 40U/µg compared to 0U/µg control). Of note, *amoA* transcripts were still quantified from the degraded 40U/µg treatment with a mean Ct of 31.8. RNase I seemed to be the most effective treatment for the destruction of rRNA. Indeed, an increase of ≈ 3.2 Cts for the *16S rRNA* transcript was observed between 0 and 40U/µg treatments whereas an increase of only 2.2 Cts was observed between 0 and 90min for both physical degradation techniques (heat and UV). R_amp_ indexes were only slightly affected by 2U RNAseI/µg (decrease of ≈0.015 for 380/120 and ≈0.03 for 380/170) (Fig. 4C.). The decrease was more pronounced for both R_amp_ at higher concentrations of RNaseI (≈0.25 decrease at 20U/µg compared to 0U control). Even at concentrations as high as 40U/µg the R_amp_ indexes only reached 0.3. This indicated that at the high nuclease concentrations, even the small amplicons (120 and 170bp) were starting to degrade. In this experiment, the R_amp_ 380/170 seemed to be more sensitive than the R_amp_ 380/120 in mapping RNA degradation, with significant differences between 0 and 10U/µg treatments whereas R_amp_ 380/120 values only became significantly different from 0U control from 20U/µg. Again, as observed in the other degradation experiments, the behaviour of the *amoA* Ct was better reflected by changes in R_amp_, especially R_amp_ 380/170, rather than by changes in the RIN.

**Fig 4.**
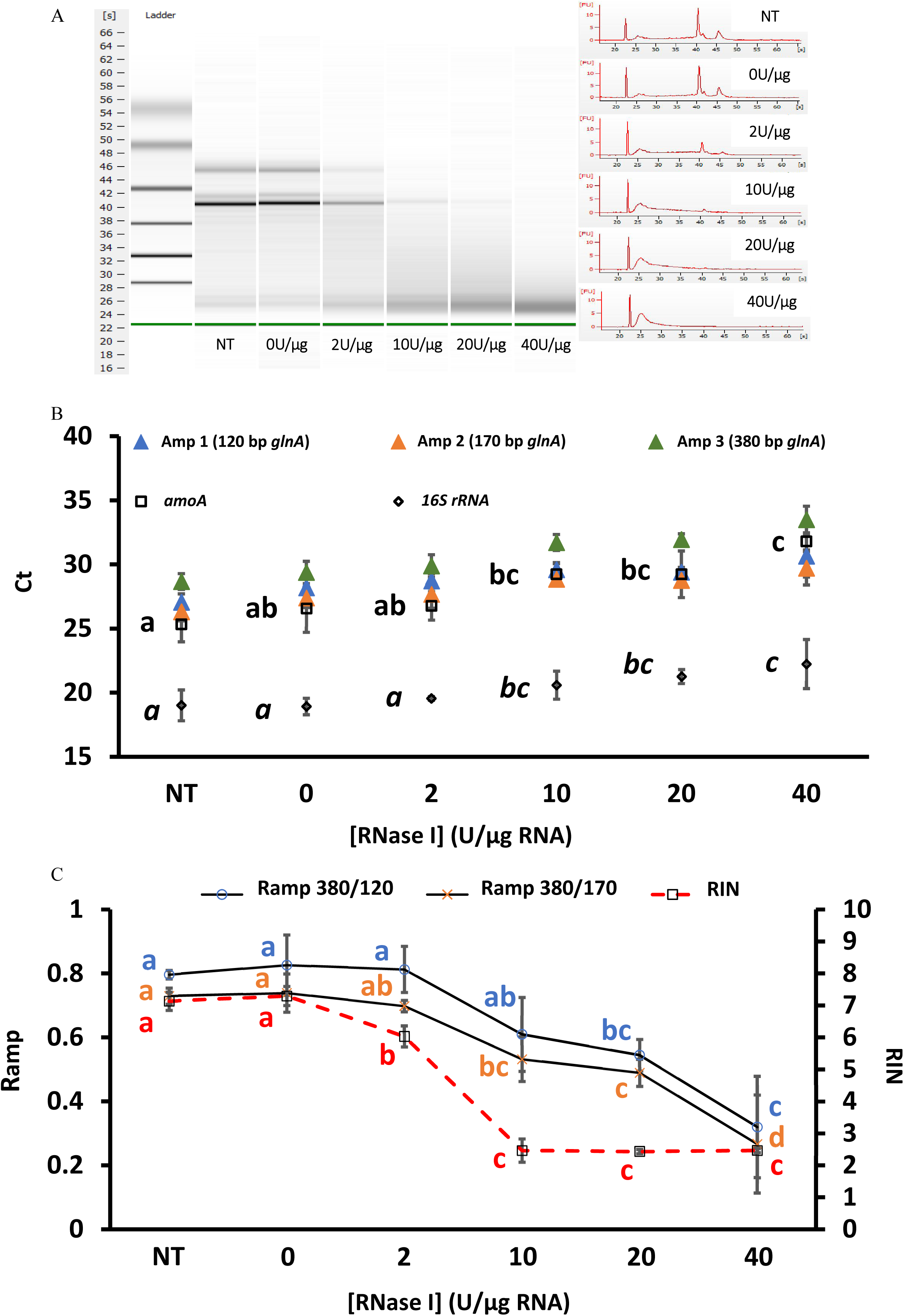
Effect of RNase I degradation on RNA integrity measured via the RIN (A), with RT-Q-PCR (B) and RIN versus R_amp_ (C). For RIN, RNA integrity visualised in virtual gels (A; left) and electropherogram (A; right) are displayed against incubation period with RNase I B) Effect of degradation on transcript quantification; Amp 1-3: average Ct (n=3) of one of the three possible *glnA* amplicons; *amoA*: average *amoA* Ct (n=3) of the Bacterial *amoA* transcript; *16S rRNA*: average *16S rRNA* Ct (n=3) of the bacterial *16S rRNA* transcript. Letters indicate the result of TukeyHSD tests (points with different letters had values significantly different from each other using 0.05 as threshold for the *p*.value). Effect of RNA degradation on R_amp_ index is presented in Fig. C. The R_amp_ 380/120 was calculated as 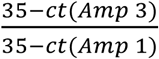 and the Ramp 380/170 as 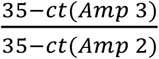; For comparison, RIN values were also plotted.

### Effect of freeze/thaw cycles

The effect of repeated cycles of freeze thaw on RNA is still poorly understood (and rarely studied) as conflicting results are reported, yet this is a common cause for concern when working with RNA. In our experiments, repeated freeze/thaw cycle (up to 10) did not induce any noticeable effects on RNA integrity, whether monitored via RIN or R_amp_ (data not shown). The effect of long term storage was also investigated but no effect could be seen after four months storage at -80°C.

### Comparison between R_amp_ and RIN

Data generated from all of the degradation experiments undertaken (UV, heat and RNaseI) was compiled to determine which of the two integrity indexes (RIN VS R_amp_) reflected the degradation status of the *amoA* and 16S rRNA transcripts more closely as determined by RT-Q-PCR. This was done by calculating Kendall correlations between either the R_amp_ or the RIN and the Cts of the two gene transcript targets (Fig 5). When considering all three degradation experiments, that is UV, heat and *RNaseI*, the RIN was not significantly correlated with 16S rRNA nor *amoA* Ct values (*p*.value > 0.05). In contrast, the R_amp_ 380/170 ratio resulted in a significant correlation with both *amoA* and 16S rRNA transcripts. The shorter R_amp_ 380/120 ratio was significantly correlated with *amoA* only (Fig 5A). However, as the RIN was almost insensitive to UV, with a decrease of only about ≈1 after 90min exposure (Fig. 2), Kendall correlations were repeated without the inclusion of the UV data set. In this case, both the RIN and the R_amp_ were significantly correlated with 16S rRNA and *amoA* transcript abundances within the degraded RNA samples (Fig. 5B). In fact, the RIN was better correlated with *amoA* than 16S rRNA Cts. Nevertheless, both R_amp_ ratios were more highly correlated with *amoA* Cts than the RIN. Furthermore, the R_amp_ approach was more highly correlated with the 16S rRNA than the RIN. Taken together, these two observations confirm that the R_amp_ indexes better reflected RT-Q-PCR changes induced by RNA degradation than the RIN.

**Fig 5.**
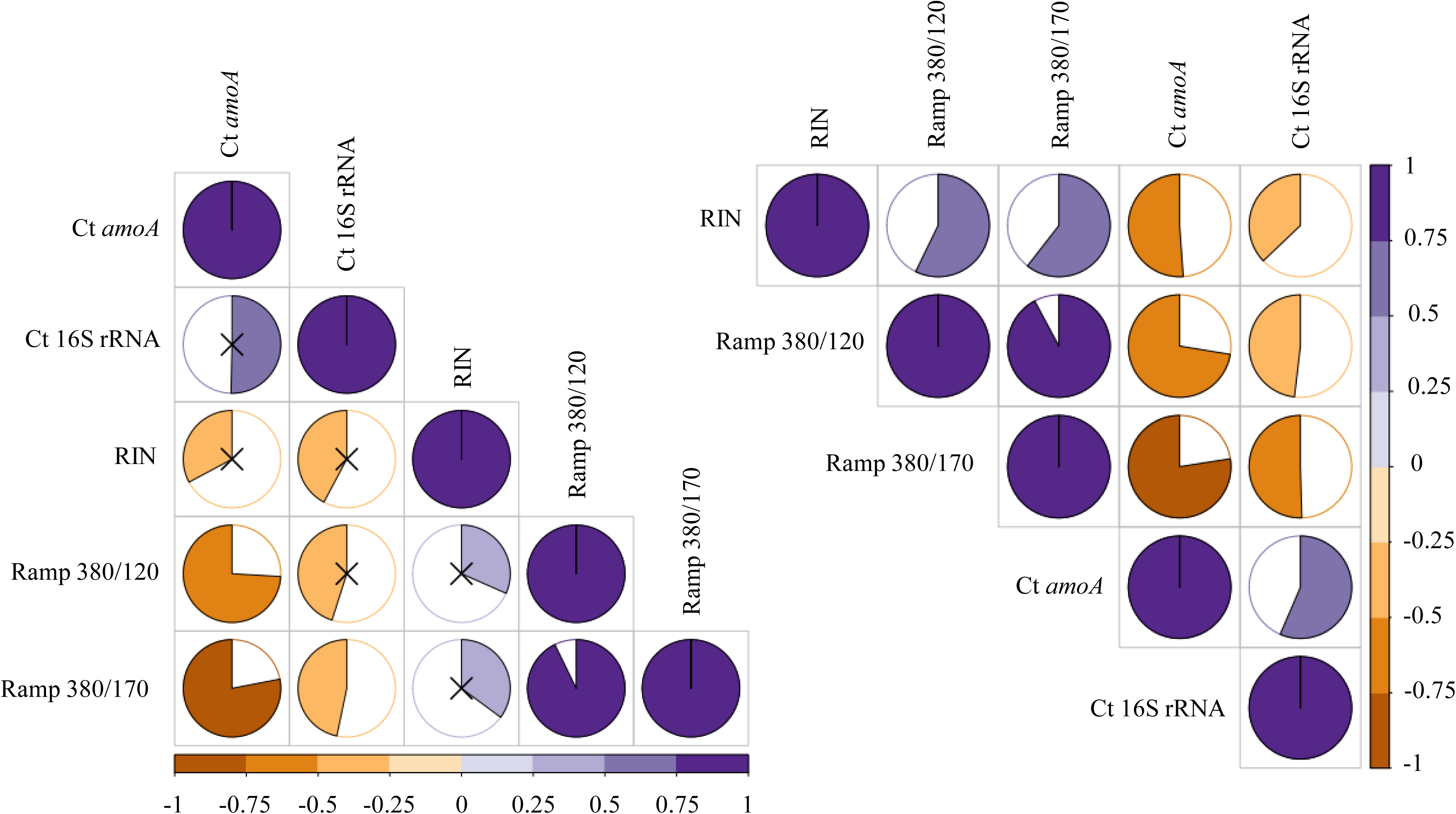
Kendall correlations between integrity indexes and Cts of the two reference gene used in this study. The correlations coefficients were calculated using all data generated from UV, heat and RNaseI degradation experiments (left) and from the heat and RNase I only (right). Black crosses indicate absence of significant correlation (threshold: p value>0.05).

### Effect of RNA degradation on transcript community composition

RNA degradation impacted upon *amoA, glnA* and *16S rRNA* gene quantification, as demonstrated previously. However, whether all members of the community were affected equally was still to be determined. To answer this question, cDNA amplicons of the Bacterial 16S rRNA, *amoA* and *glnA* transcripts underwent Illumina MISeq amplicon sequencing from all degradation points of the RNase I experiment representing RNA with RIN values from 7.5 to 2.4 and R_amp_ values from ≈0.8 to ≈0.3 and from ≈0.7 to ≈0.3 for Ramp 380/170 and Ramp 380/120 respectively. The effect of RNaseI treatment on community evenness was tested using PERMANOVA. Results are presented in Table 2 and figures 6, 7 and 8. Interestingly, the community structure of the three transcripts studied responded differently.

**Fig 6.**
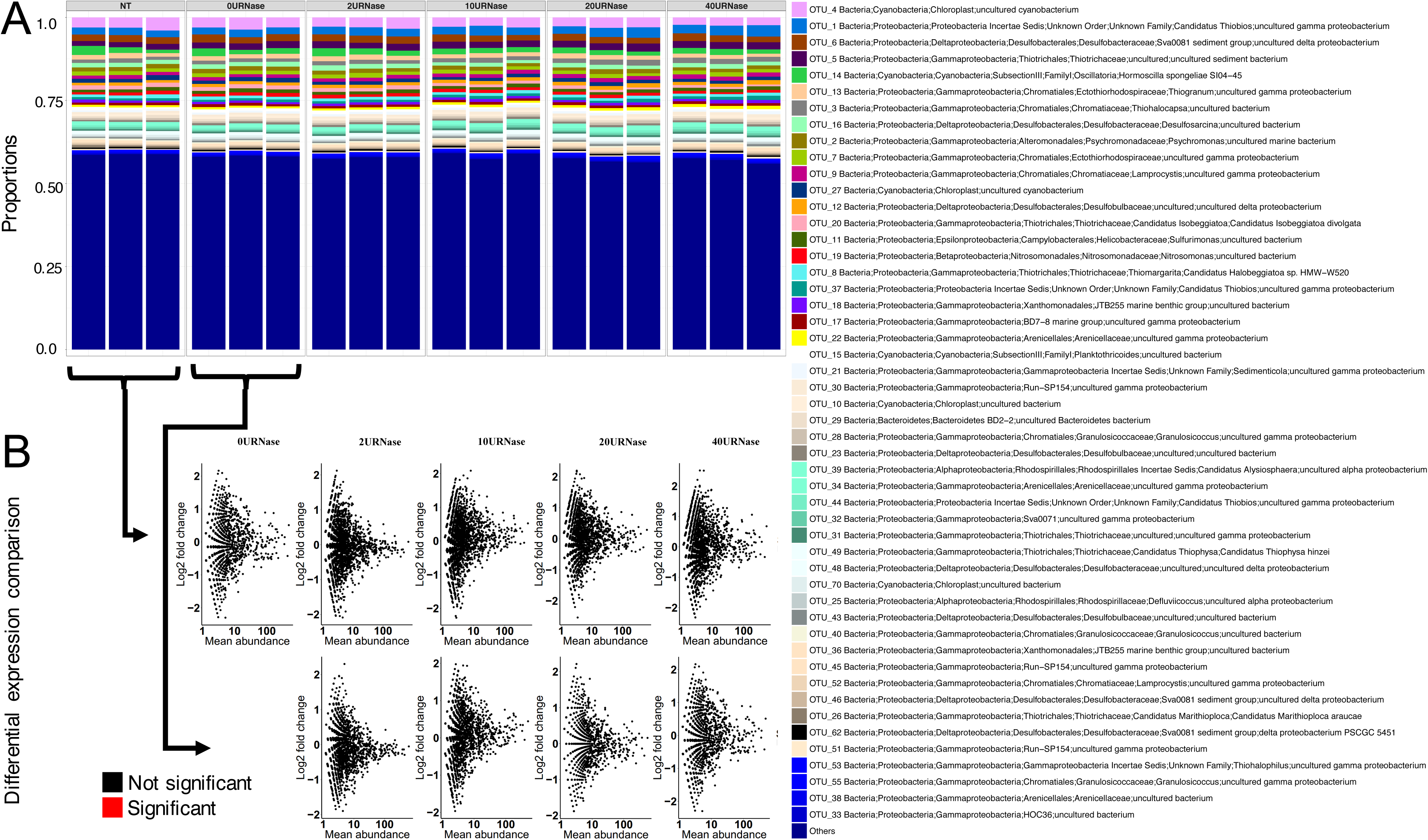
Effect of RNase I treatment on 16S rRNA transcript composition. Bar charts (A) represent changes in community composition of the 50 most abundant taxa. Scatterplots (B) represent log2 changes of individual taxa along the degradation gradient relative to control experiments (no treatment control (NT) or buffer only control (0URNaseI/µl)) as indicated by black arrows. Taxa with a significant difference (*p*.value< 0.05) in expression greater than or equal to a 2-fold change (positively or negatively) relative to controls are indicated in red.

**Fig 7.**
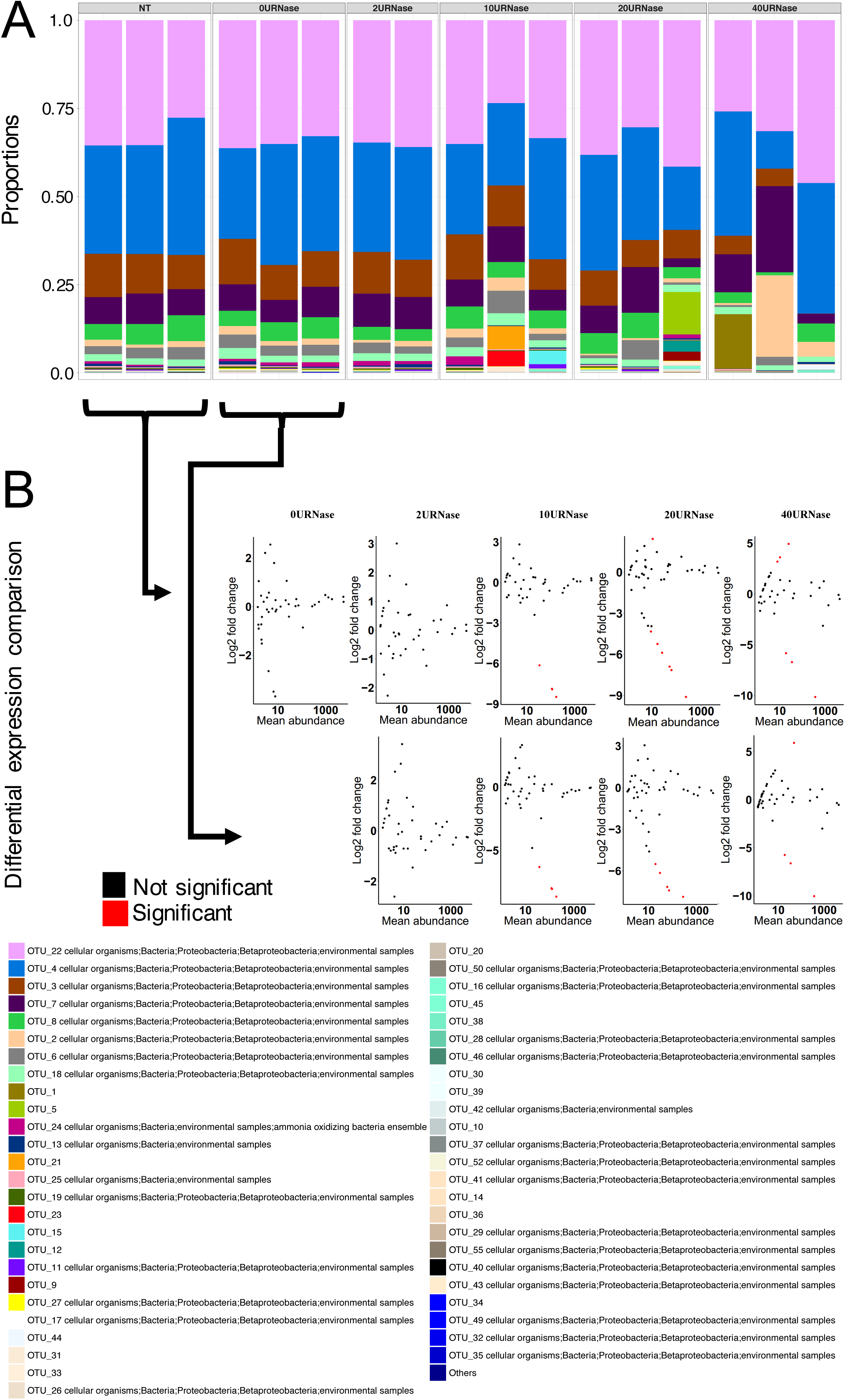
Effect of RNase I treatment on *amoA* transcript composition. Bar charts (A) represent changes in community composition of the 50 most abundant taxa. Scatterplots (B) represent log2 changes of individual taxa along the degradation gradient relative to control experiments (no treatment control (NT) or buffer only control (0URNaseI/µl)) as indicated by black arrows. Taxa with a significant difference (*p*.value< 0.05) in expression greater than or equal to a 2-fold change (positively or negatively) relative to controls are indicated in red.

**Fig 8.**
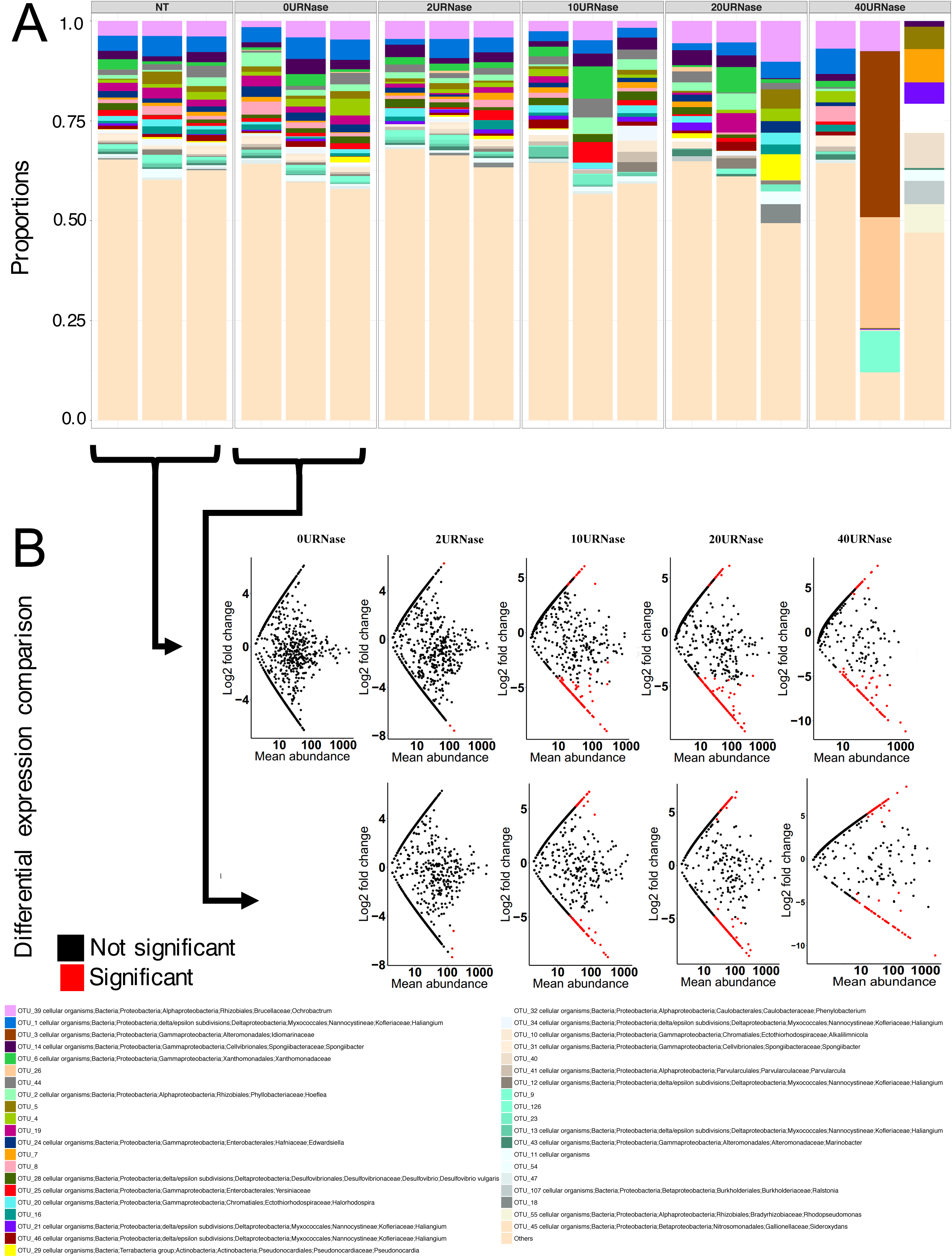
Effect of RNase I treatment on *glnA* transcript composition. Bar charts (A) represent changes in community composition of the 50 most abundant taxa. Scatterplots (B) represent log2 changes of individual taxa along the degradation gradient relative to control experiments (no treatment control (NT) or buffer only control (0URNaseI/µl)) as indicated by black arrows. Taxa with a significant difference (*p*.value< 0.05) in expression greater than or equal to a 2-fold change (positively or negatively) relative to controls are indicated in red.

Strikingly, RNase I treatment had little effect on *16S rRNA* transcript community evenness (Fig 6A). Indeed, for individual OTUs, none of the members of the community were significantly differentially represented (*p*.value log_2_ difference >0.05) within highly degraded samples in comparison to controls (Fig 6B). For individual OTU’s at least 90% had their relative expression change over the degradation experiment fall within the [-log2(1.5); lod2(1.5)] interval, even when comparing controls to the completely degraded 40U RNase I sample (Fig 6B). This indicates that *16S rRNA* OTU transcript community was responding evenly to degradation, with each member having the same chance to be affected regardless of its abundance or sequence.

For bacterial *amoA* transcript community there was no change in the overall composition with increasing degradation as reflected by the non-significant PERMANOVA (p.value >0.05). However, with increasing degradation, there was an increasing difference in the community evenness among replicates. Furthermore, unlike 16S rRNA transcripts, when examining individual *amoA* OTUs it was evident that in the degraded samples some OTUs were differentially represented at a significant level compared to controls (Fig. 7B). In fact, some OTUs in the highly degraded samples (10, 20 and 40U RNase I) had a fold change difference of up to 2 orders of magnitude compared to the controls and in most cases, resulting in their over representation in degraded samples (see Additional file 3). Moreover, in the more highly degraded treatments (10, 20 and 40U RNase I), up to 44% of *amoA* OTUs had their relative expression outside the [-log2(1.5); log2(1.5)] interval, compared to the starting RNA (Fig 7 B). So while there was not an overall significant difference in *amoA* community structure with increasing RNA degradation, there were changes in the relative expression of individual OTUs. The lack of overall statistical significance in community structure may in fact be explained by the overall lower numbers of *amoA* OTUs for comparison and the increasing difference among replicates in the degraded samples.

The effect of RNase I treatment was much more pronounced for *glnA* transcripts, than for *amoA,* and a significant change in community composition with increasing degradation was observed (*p*.value< 0.05 for PERMANOVA with both Bray-Curtis and Unifrac distances) (Fig 8A & 8B). As seen with *amoA*, the difference in community composition between replicates also increased with increasing RNase I treatment. Moreover, this effect was also observed at individual OTU level with a large fraction of the individual OTUs showing different expression levels in treated samples compared to controls (Fig. 8B). As seen for *amoA*, some *glnA* OTUs were highly over represented in degraded samples by 2 to 3 orders of magnitudes (Additional file 3), *e.g.* when comparing the untreated samples (NT) to the 40URNase samples, 0.28% (3 sequences) were over represented by 2 orders of magnitude. When comparing the samples treated with buffer only to the 40URNase samples, 2.43% (19 sequences) were over represented by 2 orders of magnitude and 0.13% (1 sequence) by 3 orders of magnitude.

## Discussion

Here we successfully designed and tested the Ratio Amplicon, Ramp, index. The concept is that as RNA degrades, longer strands are preferentially affected and the abundance of the longer amplicon relative to the shorter amplicon will decrease with increasing RNA degradation [18]. Using experimentally degraded environmental RNA we have shown that the newly developed Ramp index was a better predictor of the Ct of the target mRNA transcript used in this study, *amoA,* than the ribosome based RIN approach. In fact, when data from the three degradation experiments carried out was considered together only the Ramp statistically correlated with *amoA* Cts. As the RIN failed to detect UV degradation, we removed this data from the correlations calculation to determine if this data set was biasing the results towards the Ramp approach. In this case, there was also a significant correlation between the RIN and *amoA* Ct (−0.51). However, the Ramp index still reflected the fate of the mRNA better than the RIN (−0.72 and -0.77 for Ramp 380/120 and Ramp 380/170 respectively).

Taking the different RNA degradation approaches used individually, the RIN and Ramp ratios responded differently. As noted above, the RIN did not change over a 90-minute exposure to UV. UV causes intramolecular crosslinking of thymines but does not cause strand breaks [47] while the RIN monitors stand break. Similar results were obtained by Bjorkman *et al* [18] who reported a lack of response for the RIN and the RQI when human RNA preparations were degraded by UV radiation, even after 120 minutes of exposure. As such RNA damage by UV can’t be detected by electrophoresis separation but is recorded by RT-Q-PCR Ramp index. Other RNA degradation processes that result in base destruction but not necessarily strand break include oxidative damage [48] or chemically-induced radical formation [49].

In contrast, the RIN was the most efficient method to detect heat degradation. There was a strong and significant decrease in this index after 10 minutes whereas the Ramp indexes only became significantly different from the controls after 45 minutes. Moreover, there was very little effect on the direct quantification of the transcripts by RT-Q-PCR with very little change in the Ct of either *amoA* or 16S rRNA in the first 10 minutes at 90°C. Initially, heat degradation caused a rapid decrease in the RIN. However, at this point the RT-Q-PCR targets were actually responding more slowly and were more closely mapped by the Ramp than the RIN. Björkman *et al* [18] showed a similar response of their differential amplicon, the ΔΔamp index, that didn’t change much between 2 and 10 min at 95°C whereas the RIN rapidly reduced from 7 to 2. Moreover, Gingrich *et al* [50] showed that transcripts could be quantified from RNA preparations incubated at 90°C for several hours. This relatively low impact of heat on RNA quantification may be due to modification of RNA secondary structures which could result in more efficient cDNA synthesis and mask the effect of the heat-induced reduction of RNA integrity. More likely it is due to the small amplicon size of the targets that are unaffected by degradation. This essentially illustrates the difference in the methods used to monitor RNA degradation - the RIN detects strand break no matter where the fracture occurs along the transcript while the Ramp, will only detect degradation if the break occurs between primer binding sites.

RNA degradation using the nuclease enzyme RNase I was monitored using both RIN and Ramp. A similar behaviour could be observed here as in the heat degradation experiment with the RIN responding more quickly but loosing sensitivity when RNA was highly degraded whereas the Ramp responded slightly later but remained sensitive when RNA was extensively degraded. RNase I was the degradation method that had the strongest effect on the 16S rRNA Ct. RNase I activity is dependent on the concentration of the substrate. If rRNA and mRNA are considered as two distinct substrates, it can be expected that RNase I will have a greater impact on ribosomes as they constitute 80-85% of total RNA. Furthermore, cDNA synthesis from mRNA would be enhanced in preparations where rRNA was depleted [51]. This dynamic may mask and change the effect of degradation over time, which would explain the relatively low increase in Ct for *amoA* at the beginning of the RNase I degradation experiment. Nevertheless, in this experiment and generally, for all degradation tests carried out, the behaviour of the *amoA* Ct was better predicted by the R_amp_, as reflected by the higher correlation coefficient between R_amp_ indexes and *amoA* Ct than the RIN (Fig. 5). As the *in-vitro* half-life of different transcripts is not well understood and has been shown to vary [52–54] further work is required to test the correlation of the R_amp_ against a larger range of mRNAs. For ribosomal RNA, while the correlation between the R_amp_ index and 16S rRNA Ct was lower than for *amoA,* it still correlated better with RNA degradation than the RIN. This indicates that the outcome of 16S rRNA analysis was less affected by degradation than our mRNA targets. There are two factors that may contribute to this, the reported greater robustness of ribosomal RNA than mRNA and the shorter (∼103 bp) 16S rRNA amplicon. That ribosomal RNAs behave the same as mRNA has never been proven. On the contrary, Sidova *et al* [55] showed that when natural *post-mortem* degradation occurs, rRNA is more stable than mRNA. In this case, rRNA is a poor predictor of degradation of the mRNA fraction, as supported by this work. As mRNA is subjected to more rapid decay to adjust to the needs of the cell whereas rRNA are degraded only under certain stress conditions or when defective [56] then these intrinsic differences in stability properties may also affect degradation rates of the different class of RNA post-extraction. Therefore, based on this work we can conclude that the R_amp_ was a better predictor of mRNA integrity than the RIN.

However, as we and others [18] have shown RNA responds differently to different types of degradation e.g. strand break verses intramolecular crosslinking of thymines, and as the exact and likely multiple causes of post-extraction degradation are unknown, we recommend that the RIN is used in conjunction with the R_amp_ to monitor RNA integrity.

### Which Ramp to use?

Since, in practice, only one Ramp index is necessary, we recommend using the Ramp 380/170. In theory, the higher the difference between the two amplicons the more sensitive the index would be. We initially designed a 500bp *glnA* PCR amplicon however, the Q-PCR assay failed to produce a single diagnostic melt curve analysis. The Ramp 380/170 always had a higher value than the Ramp 380/120 which would indicate that the number of 170bp targets is higher than the 120bp. Since both are amplified from the same target, this is not possible and the explanation for this observation is the lower efficiency of the 120bp Q-PCR compared to the 170bp assay. In spite of this, both Ramp correlated similarly well overall with each degradation experiment, with Ramp 380/170 slightly more sensitive in the RNase I experiment.

### Impact of experimental degradation of environmental RNA on ribosomal (16S rRNA) and mRNA (*amoA*) community diversity

For complex environmental communities, the integrity of RNA is not only important to evaluate quantitative gene expression, but is also of significance if it adversely affects the relative abundance of transcript diversity. To examine this, we assessed changes in the community structure of the 16S rRNA, *amoA* and *glnA* transcripts from all fractions of the RNase I sequentially degraded RNA.

The results were surprising with successful amplicon sequencing even from highly degraded samples. Nevertheless, the data did suggest a different response of 16S rRNA and mRNA transcripts to degradation, with 16S rRNA community structure unaffected over the range of degraded RNA samples. That is a statistically similar community was present in the control non-degraded samples as in the totally destroyed 40 Units RNase I (with a mean RIN of 2.5 and Ramp of 0.32 and 0.27 for Ramp 380/120 and Ramp 38/170 respectively). This indicates that while total RNA was degraded, the small transcript fragments required for RT-PCR and amplicon sequencing remained intact. In fact, so much so that no significant change in the relative abundance of individual OTUs was observed.

On the other hand, RNA degradation had a greater influence on both *amoA* and *glnA* mRNA targets. While, again surprisingly, transcript amplicons were successfully detected from all degradation status samples, greater variability between degraded replicates was observed. This resulted in statistically different communities for *glnA* but not *amoA* when compared to the same non-degraded control samples. However, the low number of *amoA* OTUs and increased variability between replicates contributed to the lower statistical power resulting in no statistical difference between treatments (Fig. 6). Furthermore, there were significant, sometimes up to 2 to 3 orders of magnitude change in the relative abundance of individual *glnA* and *amoA* OTUs in the degraded samples verses control samples. So, while we could successfully amplify mRNA transcripts from degraded environmental samples, we have shown that the relative composition of the community members was adversely affected by degradation and was not representative of the initial starting point. While further work is needed to determine the impact of degradation across the entire transcriptome to see if all mRNA’s respond in a similar manner, it is clear from our mRNA amplicon sequencing that RNA degradation will alter the outcome of community analysis. It is therefore necessary to ensure the RNA integrity of the sample is known prior to interpretation of results. For this our data indicates that a combination approach targeting both ribosomal (the RIN) and mRNA (the Ramp) is needed.

### Best Practice for Environmental RNA

The challenge when working with environmental samples will always be to retrieve RNA of a high enough quality and integrity. Here we started with RNA extracted from marine sediments that had an average RIN of ≈7 and R_amp_ of ≈0.8. This is the best quality RNA we could produce with this beat-beating co-extraction method [36] and it already falls at the lower end of acceptable RIN for pure culture [10]. Therefore, methods to improve the initial quality of RNA extractions should also be a high priority, although this will be easier in some environments than others. Improvement of extraction methods is crucial as it can lead to important differences in the results. For example Feike *et al* [57] showed that different sampling techniques influenced the relative abundance of transcripts retrieved from the suboxic zone of the Baltic Sea. Next, the integrity of the extracted RNA should be determined, and it should be ensured that the integrity value is similar among samples to be compared. Here the R_amp_ approach should be a useful tool to complement current electrophoretic approaches, such as the RIN prior to extensive downstream analysis.

Another consideration raised by this work is in the very fact that the differential amplicon approach works. This shows that small cDNA amplicons can still be produced from highly degraded RNA samples whereas long amplicons tend to disappear quickly. When using RNA samples of poor quality, the comparison of expression levels between different targets might be irrelevant if the difference in length of the RT-Q-PCR targets between genes is large. In this case it would be better to use only small amplicons, that are less sensitive to degradation [58]. An alternative, to deal with samples with different degradation status, potentially could be to normalize RT-Q-PCR data to RNA integrity. A RIN based algorithm has been proposed by Ho-Pun-Cheung *et al* [59] to reduce RT-Q-PCR errors due to RNA degradation in cancer biopsies. In our case however, Ramp indexes correlated better than the RIN with *amoA* and 16S rRNA Cts, making them better potential candidates as normalisation metrics. Therefore we tested a normalisation coefficient based on the Ramp (Additional file 2; Figure S2). As in Ho-Pun-Cheung *et al* [59], we assumed a linear relationship between the integrity index and the changes in transcript Cts (*i.e.* change in Ct = α x change in Ramp). This assumption facilitated the calculation of a regression coefficient α that was used to normalize Cts as explained in figure S2. Although the use of such normalization reduced the errors attributable to RNA degradation (Additional file 2; Figure S2.), several limitations remain: 1) the linear relationship between changes in Cts and Ramp might not always be true depending on the transcript tested, 2) the regression coefficient α depends on the degradation technique (Additional file 2; Table S1), 3) the regression coefficient α depends on the transcript tested (Additional file 2; Table S1) and 4) the regression coefficient α may depend on the environment from which RNA was extracted. Until more work is done to validate such normalization strategies, or to dramatically improve the quality of the RNA that can be extracted from environmental samples [57], we recommend using integrity indexes (differential amplicon and microfluidics based techniques) as initial quality checks of RNA and advise not to make absolute comparisons among samples with dissimilar integrity status.

## Conclusion

Assessing RNA quality is essential for obtaining meaningful transcriptomic results. The current approach to monitor RNA integrity include the RIN and RQI. This is a useful technique that is widely under-used (or reported) in microbial transcriptomics studies, to give an overview of total RNA quality based on a ratio between the 23S and 16S ribosomes. Since most transcriptomics studies are interested in the metabolic function and therefore mRNA, it would be preferable to have an integrity index to target the mRNA. Furthermore, it is unknown if degradation of rRNA reflects mRNA degradation. We therefore developed and experimentally tested a new index, the Ramp, the goal of which was to specifically target mRNA degradation and we showed that it performed better than the RIN at predicting the outcome of RT-Q-PCR of a functional gene (*amoA*). It was shown in this study that both quantitative (RT-Q-PCR) and qualitative (sequencing) results can be obtained, even from very degraded samples. Comparison of gene expression level between preparations with different degradation levels can therefore lead to false conclusions if integrity is not checked prior to analysis. Thus, we encourage microbial ecologists to report integrity indexes in order to improve reproducibility and facilitate comparison between transcriptomics studies. For this we propose that a Ramp ratio is used alongside the RIN.

## Abbreviations

R_amp_: Ratio Amplicon
RIN: RNA Integrity number
RQI: RNA Quality Score
RT-(Q)-PCR: Reverse Transcriptase (Quantitative) Polymerase Chain Reaction
cDNA: complementary DNA
Ct: Cycle threshold

## Ethics approval and consent to participate

Not applicable

## Consent for publication

Not applicable

## Authors’ contribution

CJS supervised the project. CJS and FC designed the experiments. FC carried out the experiments. FC and CJS analysed the RT-Q-PCR and RIN measurment data. FC and UZI ran the bioinformatic pipeline for amplicon sequences processing. FC, CJS, UZI and analysed the amplicon sequencing results. FC, UZI, CJS and wrote the manuscript. All authors read and approved the final manuscript.

## Data availability

Ct and RIN data presented in this study are provided in additionnal file 7. The sequencing data are available on the European Nucleotide Archive under the study accession number: PRJEB28215 (http://www.ebi.ac.uk/ena/data/view/PRJEB28215) with information about the samples given in additional file 5 and 6.

## Acknowledgements

We would like to thank Dr. Aoife Duff for collecting the sediment samples.

## References

1. Medini D, Serruto D, Parkhill J, Relman DA, Donati C, Moxon R, et al. Microbiology in the post-genomic era. Nat Rev Microbiol. 2008;6:419–30.

2. Moran MA, Satinsky B, Gifford SM, Luo H, Rivers A, Chan LK, et al. Sizing up metatranscriptomics. ISME J. 2013;7:237–43. doi:10.1038/ismej.2012.94.

3. Evans TG. Considerations for the use of transcriptomics in identifying the “genes that matter” for environmental adaptation. J Exp Biol. 2015;218:1925–35. doi:10.1242/jeb.114306.

4. Smith CJ, Nedwell DB, Dong LF, Osborn AM. Evaluation of quantitative polymerase chain reaction-based approaches for determining gene copy and gene transcript numbers in environmental samples. Environ Microbiol. 2006;8:804–15.

5. Laalami S, Zig L, Putzer H. Initiation of mRNA decay in bacteria. Cell Mol Life Sci. 2014;71:1799–828.

6. Die J V, Román B. RNA quality assessment: a view from plant qPCR studies. J Exp Bot. 2012;63:6069–77.

7. Copois V, Bibeau F, Bascoul-Mollevi C, Salvetat N, Chalbos P, Bareil C, et al. Impact of RNA degradation on gene expression profiles: Assessment of different methods to reliably determine RNA quality. J Biotechnol. 2007;127:549–59.

8. Fleige S, Pfaffl MW. RNA integrity and the effect on the real-time qRT-PCR performance. Mol Aspects Med. 2006;27:126–39.

9. Fleige S, Walf V, Huch S, Prgomet C, Sehm J, Pfaffl MW. Comparison of relative mRNA quantification models and the impact of RNA integrity in quantitative real-time RT-PCR. Biotechnol Lett. 2006;28:1601–13.

10. Jahn CE, Charkowski AO, Willis DK. Evaluation of isolation methods and RNA integrity for bacterial RNA quantitation. J Microbiol Methods. 2008;75:318–24.

11. Kaczanowska M, Ryden-Aulin M. Ribosome Biogenesis and the Translation Process in Escherichia coli. Microbiol Mol Biol Rev. 2007;71:477–94. doi:10.1128/MMBR.00013-07.

12. Rhodes ME, Oren A, House CH. Dynamics and persistence of dead sea microbial populations as shown by high-throughput sequencing of rRNA. Appl Environ Microbiol. 2012;78:2489–92.

13. McKillip JL, Jaykus L, Drake M. rRNA Stability in Heat-Killed and UV-Irradiated Enterotoxigenic Staphylococcus aureus and Escherichia coli O157?: H7 rRNA Stability in Heat-Killed and UV-Irradiated Enterotoxigenic Staphylococcus aureus and Escherichia coli O157?: H7. Appl Environ Microbiol. 1998;64:4264–8.

14. Rathnayaka USK, Rakshit SK. The stability of rrna in heat-killed Salmonella enterica cells and its detection by fluorescent in situ hybridisation (FISH). Trop Life Sci Res. 2010;21:47–53.

15. Hurt RA, Qiu X, Wu L, Roh Y, Palumbo A V, Tiedje JM, et al. Simultaneous Recovery of RNA and DNA from Soils and Sediments. Appl Environ Microbiol. 2001;67:4495–503.

16. Die J V., Obrero Á, González-Verdejo CI, Román B. Characterization of the 3’:5’ ratio for reliable determination of RNA quality. Anal Biochem. 2011;419:336–8. doi:10.1016/j.ab.2011.08.012.

17. Dreyfus M, Régnier P. The poly(A) tail of mRNAs: Bodyguard in eukaryotes, scavenger in bacteria. Cell. 2002;111:611–3.

18. Björkman J, Švec D, Lott E, Kubista M, Sjöback R. Differential amplicons (δAmp)-a new molecular method to assess RNA integrity. Biomol Detect Quantif. 2016;6:4–12.

19. Karlsson O, Segerström L, Sjöback R, Nylander I, Borén M. qPCR based mRNA quality score show intact mRNA after heat stabilization. Biomol Detect Quantif. 2016;7:21–6. doi:10.1016/j.bdq.2016.01.002.

20. Reck M, Tomasch J, Deng Z, Jarek M, Husemann P, Wagner-Döbler I. Stool metatranscriptomics: A technical guideline for mRNA stabilisation and isolation. BMC Genomics. 2015;16:494. doi:10.1186/s12864-015-1694-y.

21. Kumada Y, Benson DR, Hillemann D, Hosted TJ, Rochefort D a, Thompson CJ, et al. Evolution of the glutamine synthetase gene, one of the oldest existing and functioning genes. Proc Natl Acad Sci U S A. 1993;90:3009–13.

22. Brown JR, Masuchi Y, Robb FT, Doolittlel WF. Evolutionary relationships of bacterial and archaeal glutamine synthetase genes. J Mol Evol. 1994;38:566–76.

23. Reitzer L. Nitrogen Assimilation and Global Regulation in *Escherichia coli*. Annu Rev Microbiol. 2003;57:155–76. doi:10.1146/annurev.micro.57.030502.090820.

24. Costa R, Gomes NCM, Milling A, Smalla K. An optimized protocol for simultaneous extraction of DNA and RNA from soils. Brazilian J Microbiol. 2004;35:230–4.

25. Sharma S, Mehta R, Gupta R, Schloter M. Improved protocol for the extraction of bacterial mRNA from soils. J Microbiol Methods. 2012;91:62–4. doi:10.1016/j.mimet.2012.07.016.

26. Sessitsch A, Gyamfi S, Stralis-Pavese N, Weilharter A, Pfeifer U. RNA isolation from soil for bacterial community and functional analysis: evaluation of different extraction and soil conservation protocols. J Microbiol Methods. 2002;51:171–9. doi:S0167701202000659 [pii].

27. Atkinson MR, Blauwkamp T a, Bondarenko V, Studitsky V, Ninfa AJ. Activation of the glnA, glnK, and nac Promoters as Escherichia coli Undergoes the Transition from Nitrogen Excess Growth to Nitrogen Starvation. Society. 2002;184:5358–63.

28. Hua Q, Yang C, Oshima T, Mori H, Shimizu K. Analysis of Gene Expression in Escherichia coli in Response to Changes of Growth-Limiting Nutrient in Chemostat Cultures. Society. 2004;70:2354–66. doi:10.1128/AEM.70.4.2354.

29. Leigh JA, Dodsworth JA. Nitrogen Regulation in Bacteria and Archaea. Annu Rev Microbiol. 2007;61:349–77. doi:10.1146/annurev.micro.61.080706.093409.

30. Duff AM, Zhang LM, Smith CJ. Small-scale variation of ammonia oxidisers within intertidal sediments dominated by ammonia-oxidising bacteria Nitrosomonas sp. amoA genes and transcripts. Sci Rep. 2017;7:1–13. doi:10.1038/s41598-017-13583-x.

31. Zhang L, Duff A, Smith C. Community and functional shifts in ammonia oxidizers across terrestrial and marine (soil/sediment) boundaries in two coastal Bay ecosystems. Environ Microbiol. 2018;00.

32. Clark K, Karsch-Mizrachi I, Lipman DJ, Ostell J, Sayers EW. GenBank. Nucleic Acids Res. 2016;44:D67–72.

33. Altschul SF, Gish W, Miller W, Myers EW, Lipman DJ. Basic local alignment search tool. J Mol Biol. 1990;215:403–10. doi:10.1016/S0022-2836(05)80360-2.

34. Edgar RC. MUSCLE: Multiple sequence alignment with high accuracy and high throughput. Nucleic Acids Res. 2004;32:1792–7.

35. Kumar S, Stecher G, Tamura K. MEGA7: Molecular Evolutionary Genetics Analysis Version 7.0 for Bigger Datasets. Mol Biol Evol. 2016;33:1870–4.

36. Griffiths RI, Whiteley AS, O’Donnell AG, Bailey MJ. Rapid method for coextraction of DNA and RNA from natural environments for analysis of ribosomal DNA- and rRNA-based microbial community composition. Appl Environ Microbiol. 2000;66:5488–91.

37. Winter DJ. rentrez: An R package for the NCBI eUtils API. R J. 2017;9:520–6. doi:10.7287/peerj.preprints.3179v2.

38. Joshi NA, Sickle JNF. A sliding-window, adaptive, quality-based trimming tool for FastQ files. 2011;:1–9.

39. Nikolenko SI, Korobeynikov AI, Alekseyev MA. BayesHammer: Bayesian clustering for error correction in single-cell sequencing. BMC Genomics. 2013;14 Suppl 1:1–11.

40. Schirmer M, Ijaz UZ, D’Amore R, Hall N, Sloan WT, Quince C. Insight into biases and sequencing errors for amplicon sequencing with the Illumina MiSeq platform. Nucleic Acids Res. 2015;43.

41. D’Amore R, Ijaz UZ, Schirmer M, Kenny JG, Gregory R, Darby AC, et al. A comprehensive benchmarking study of protocols and sequencing platforms for 16S rRNA community profiling. BMC Genomics. 2016;17. doi:10.1186/s12864-015-2194-9.

42. Caporaso JG, Kuczynski J, Stombaugh J, Bittinger K, Bushman FD, Costello EK, et al. QIIME allows analysis of high-throughput community sequencing data. Nat Methods. 2010;7:335–6. doi:10.1038/nmeth.f.303.QIIME.

43. Katoh K, Asimenos G, Toh H. Multiple Alignment of DNA Sequences with MAFFT. In: Bioinformatics for DNA Sequence Analysis, Methods in Molecular Biology. 2009. p. 39–64. doi:10.1007/978-1-59745-251-9.

44. Price MN, Dehal PS, Arkin AP. FastTree 2 - Approximately maximum-likelihood trees for large alignments. PLoS One. 2010;5.

45. Oksanen J, Kindt R, O’Hara RB. vegan: Community Ecology Package. Available from http://cc.oulu.fi/∼jarioksa/. 2005; 3 January. http://cc.oulu.fi/∼jarioksa/.

46. Love MI, Huber W, Anders S. Moderated estimation of fold change and dispersion for RNA-seq data with DESeq2. Genome Biol. 2014;15:1–21.

47. Kladwang W, Hum J, Das R. Ultraviolet Shadowing of RNA Can Cause Significant Chemical Damage in Seconds. Sci Rep. 2012;2:517. doi:10.1038/srep00517.

48. Rhee Y, Valentine MR, Termini J. Oxidative base damage in RNA detected by reverse transcriptase. Nucleic Acids Res. 1995;23:3275–82.

49. Hawkins CL, Davies MJ. Hypochlorite-induced damage to DNA, RNA, and polynucleotides: Formation of chloramines and nitrogen-centered radicals. Chem Res Toxicol. 2002;15:83–92.

50. Gingrich J, Rubio T, Karlak C. Effect of RNA degradation on the data quality in quantitative PCR and microarray experiments. Bio-Rad Bull. 2008;:1–6. http://www.gmo-qpcr-analysis.info/gingrich-bulletin-5547A.pdf.

51. Petrova OE, Garcia-Alcalde F, Zampaloni C, Sauer K. Comparative evaluation of rRNA depletion procedures for the improved analysis of bacterial biofilm and mixed pathogen culture transcriptomes. Sci Rep. 2017; 7 May 2016:41114. doi:10.1038/srep41114.

52. Belasco JG. All Things Must Pass: Constrasts and Commonalities in Eukaryotic and Bacterial mRNA Decay. Nat Rev Mol Cell Biol. 2010;11:467–478.

53. Evguenieva-Hackenberg E, Klug G. New aspects of RNA processing in prokaryotes. Curr Opin Microbiol. 2011;14:587–92. doi:10.1016/j.mib.2011.07.025.

54. Selinger WD, Saxena MR, Cheung K., Church MG, Rosenow C. Global RNA Half-Life Analysis in. Genome Res. 2003; January:216–23.

55. Sidova M, Tomankova S, Abaffy P, Kubista M, Sindelka R. Effects of post-mortem and physical degradation on RNA integrity and quality. Biomol Detect Quantif. 2015;5:3–9. doi:10.1016/j.bdq.2015.08.002.

56. Deutscher MP. Degradation of RNA in bacteria: Comparison of mRNA and stable RNA. Nucleic Acids Res. 2006;34:659–66.

57. Feike J, Jürgens K, Hollibaugh JT, Krüger S, Jost G, Labrenz M. Measuring unbiased metatranscriptomics in suboxic waters of the central Baltic Sea using a new in situ fixation system. ISME J. 2012;6:461–70.

58. Antonov J, Goldstein DR, Oberli A, Baltzer A, Pirotta M, Fleischmann A, et al. Reliable gene expression measurements from degraded RNA by quantitative real-time PCR depend on short amplicons and a proper normalization. Lab Invest. 2005;85:1040–50.

59. Ho-Pun-Cheung A, Bascoul-Mollevi C, Assenat E, Boissière-Michot F, Bibeau F, Cellier D, et al. Reverse transcription-quantitative polymerase chain reaction: Description of a RIN-based algorithm for accurate data normalization. BMC Mol Biol. 2009;10:1–10.

60. Hornek R, Pommerening-Röser A, Koops HP, Farnleitner AH, Kreuzinger N, Kirschner A, et al. Primers containing universal bases reduce multiple amoA gene specific DGGE band patterns when analysing the diversity of beta-ammonia oxidizers in the environment. J Microbiol Methods. 2006;66:147–55.

61. Suzuki MT, Taylor LT, Delong EF, Long EFDE. Quantitative Analysis of Small-Subunit rRNA Genes in Mixed Microbial Populations via 5 ′ -Nuclease Assays Quantitative Analysis of Small-Subunit rRNA Genes in Mixed Microbial Populations via 5 Ј -Nuclease Assays. 2000;66:4605–14.

62. Marchesi JR, Sato T, Weightman AJ, Martin TA, Fry JC, Hiom SJ, et al. Design and evaluation of useful bacterium-specific PCR primers that amplify genes coding for bacterial 16S rRNA. Appl Environ Microbiol. 1998;64:795–9.

63. Lee TK, Van Doan T, Yoo K, Choi S, Kim C, Park J. Discovery of commonly existing anode biofilm microbes in two different wastewater treatment MFCs using FLX Titanium pyrosequencing. Appl Microbiol Biotechnol. 2010;87:2335–43.

64. Parada AE, Needham DM, Fuhrman JA. Every base matters: Assessing small subunit rRNA primers for marine microbiomes with mock communities, time series and global field samples. Environ Microbiol. 2016;18:1403–14.

65. Caporaso JG, Kuczynski J, Stombaugh J, Bittinger K, Bushman FD, Costello EK, et al. correspondence QIIME allows analysis of high-throughput community sequencing data Intensity normalization improves color calling in SOLiD sequencing. Nat Publ Gr. 2010;7:335–6. doi:10.1038/nmeth0510-335.

